# Modelling neural entrainment and its persistence: influence of frequency of stimulation and phase at the stimulus offset

**DOI:** 10.1101/2021.09.10.459802

**Authors:** Mónica Otero, Caroline Lea-Carnall, Pavel Prado, María-José Escobar, Wael El-Deredy

## Abstract

The entrainment (synchronization) of brain oscillations to the frequency of sensory stimuli is a key mechanism that shapes perceptual and cognitive processes, such that atypical neural entrainment leads to neuro-psychological deficits.

**Objective:** We investigated the dynamic of neural entrainment. Particular attention was paid to the oscillatory behavior that succeed the end of the stimulation, since the persistence (reverberation) of neural entrainment may condition future sensory representations based on predictions about stimulus rhythmicity.

**Approach:** A modified Jansen-Rit neural mass model of coupled cortical columns generated a time series whose frequency spectrum resembled that of the electroencephalogram. We evaluated spectro-temporal features of entrainment, during and after rhythmic stimulation of different frequencies, as a function of the resonance frequency of the neural population and the coupling strength between cortical columns. We tested if the duration of the entrainment persistence depended on the state of the neural network at the time the stimulus ends.

**Main Results:** The entrainment of the column that received the stimulation was maximum when the frequency of the entrainer was within a narrow range around the resonance frequency of the column. When this occurred, entrainment persisted for several cycles after the stimulus terminated, and the propagation of the entrainment to other columns was facilitated. Propagation depended on the resonance frequency of the second column, and the coupling strength between columns. The duration of the persistence of the entrainment depended on the phase of the neural oscillation at the time the entrainer terminated, such that falling phases (from *π*/2 to 3*π*/2 in a sine function) led to longer persistence than rising phases (from 0 to *π*/2 and 3*π*/2 to 2*π*).

**Significance:** The study bridges between models of neural oscillations and empirical electrophysiology, and provides insights to the use of rhythmic sensory stimulation for neuroenhancement.

## 1. Introduction

Sensory stimulation has being used for the modulation of brain oscillations for clinical and non-clinical purposes due to their safety and effectiveness [1–5]. As a matter of fact, neuromodulation is considered a promising therapeutic tool that promotes cognitive enhancement by increasing the ability of distributed cortical networks to coordinate and generates brain rhythms. The impairment of this oscillatory dynamics, which is referred to as oscillophaties, can result in altered behavioral outcomes, and has impact on several neurological and psychiatric disorders including schizophrenia, Alzheimer disease, Parkinson’s disease, epilepsy and sleep disorders [6–13].

The theoretical framework supporting neuromodulation states that the stimulus-driving neural oscillations result from the synchronization (or coupling) of neural oscillations to the frequency of external stimuli [14–17]. This process is referred to as neural entrainment, and is argued to be a basic mechanism that shapes sensory perception. Neural entrainment is crucial for structuring incoming information streams for further processing, including, attention selection, learning, and motor execution [14–18]. Speech perception, and music appreciation in particular, rely on neural entrainment to extract relevant features from the continuous acoustic signals [14,19]. When endogenous brain oscillations phase align to salient events in the sound stream, particularly in the delta/theta (1-7.5 Hz) frequency band of the electroencephalogram (EEG), the events are processed and perceived more readily than temporally non-overlapping events [20–22].

Although well characterized from a phenomenological perspective, neuro-modulation therapies based on periodic brain stimulation generally lacks mechanistic models of how sensory stimulation interacts with the underlying brain oscillations, causing changes in behavioral and perceptual responses [23]. These neural stimulation tools may benefit from mathematical models of entrainment that contain plausible physiological approximations.

Entrainment arises due to the phase realignment of endogenous oscillations to the driving stimulus [14–17,24]. When multiple oscillators are simultaneously recorded in a mean field activity as in the case of EEG, their aligned phases add up, leading to an increase in amplitude of the EEG oscillation [25]. Consequently, when periodic sensory stimulation is presented, entrainment can lead to the generation of a steady-state responses, i.e., scalp-recorded brain oscillations locked to the periodicity of the sensory input. Typically, the steady-state responses are observed as an increase in the power spectrum at the frequency of the driving stimulus [26].

Empirical studies have found that the amplitude of entrained oscillations does not return to the baseline immediately after the stimulus offset but remains relatively high for approximately three consecutive cycles, i.e., the entrainment *persists* after the stimulus offset [24, 27–33]. Consistent evidence supports the idea that the persistence of this stimulus-driven brain oscillations after the stimulus offset represents a coding mechanism of temporal expectations [27,34]. In the visual domain, the persistence of alpha EEG oscillations elicited by sinusoidally-varying light reflects the functioning of fronto-occipital neuronal circuits, which may serve to prime the sensory representation of incoming visual stimuli based on predictions about stimulus rhythmicity [27]. Furthermore, the persistence of the EEG alpha entrainment depends on the phase of the stimulus-driven oscillatory activity at the time the visual stimulus is removed. While longer persistence duration is observed when the visual stimulation terminates towards the troughs of the alpha oscillations, the persistence of the entrainment is shortened when the stimuli terminate near the peaks of the EEG oscillation [27].

There are no concrete theories describing how the entrainment persists after the sensory stimulus is removed, or how the entrained signal is propagated between neural regions. Additionally, the functional relationship between the frequency of the driving stimulus, the intrinsic oscillatory properties of the neural circuit involved, and the duration of the persistence is not well understood. While addressing this open question experimentally is challenging, mathematical modelling provides a robust approach to investigating mechanisms of neural entrainment using biophysical, generative models of EEG oscillations, such as the modified version of the Jansen-Rit neural mass model proposed by [35].

The Jansen-Rit NMM is a biophysical representation of the average activity of a neural assembly, which can be thought of as a cortical column or even a cortical region [36–38]. In its simplest form, Jansen-Rit NMM is able to reproduce spectral features observed in EEG recordings but is limited to generating activity within a single narrow frequency band around 10 Hz (alpha) [38–40]. This model generated oscillatory EEG-like signals with a particular preferred oscillatory frequency, (referred as to the resonance frequency), which correspond to the frequency at which a peak in the power spectrum of the EEG is obtained. In the model, the resonance frequency of the oscillation is mostly determined by the parameters controlling the the postsynaptic membrane potential.

In order to simulate the full EEG spectrum, extensions to the original Jansen-Rit NMM have been established. In this paper we focus on the work by David and Friston [35] who proposed a neural mass consisting of multiple neural populations, such that each population generates activity at a particular resonance frequency. The output of the model is a weighted sum of each of these populations allowing the simulation of rich dynamics much closer to that of physiological EEG recordings.Therefore, a preferred oscillatory frequency emerged in the simulated EEG-like oscillation, being determined by the relative proportion of neurons of the NMM tuned to a particular frequency.

In this study we employed a Jansen-Rit NMM comprising two coupled cortical columns, to represent the entrainment of intrinsic oscillations in the EEG alpha band elicited by a sinusoidally varying light. We investigated possible mechanisms that sustain the neural entrainment, as well as the persistence of the stimulus-driven oscillation after the end of the stimulation. We analyzed the dynamics of the entrainment as a function of the match between the resonance frequency of the NMM and the frequency of the driving force. Furthermore, we investigated the relation between the duration of the persistence of the entrainment and the terminating phase of the oscillatory EEG-like activity of the NMM at the stimulus offset. Since it is well known that primary visual cortex is hierarchically connected with other cortical regions processing visual information, and that previous investigations have examined the propagation of oscillatory signals between cortical columns [35, 38], we analyzed possible physiological factors affecting the propagation of the entrainment. Our results contributes to understand neural entrainment. More remarkable still, the results can contribute to enhance the specificity and effectiveness of established neural stimulation tools, therefore impacting the design of stimulation strategies for neuro-modulation therapies based on periodic brain stimulation.

## 2. Methods

### 2.1. Model of entrainment using a Jansen-Rit NMM approach

Here we use a modified version of the Jansen-Rit NMM, proposed by David and Friston [35] which allowed us to account for two populations; one tuned to alpha and the other to gamma band resonance frequencies 1). The model is able to produce richer dynamics than the standard Jansen-Rit model allowing for generation of the full EEG spectrum.

We drove the model with ‘external’ repetitive stimulus at a range of frequencies to study the mechanisms of the entrainment and its persistence. We connected two similar cortical columns, coupled in cascade (i.e., unidirectionally from column 1 to column 2), following the approach in [35, 38] to investigate the conditions under which the entrainment propagates from one unit to the next (Figure 2).

#### 2.1.1. Double population Jansen-Rit NMM column

In the model, it is assumed that a particular cortical area comprises several resonant neuronal circuits, where each circuit is tuned to a particular frequency. This idea is supported by experimental results demonstrating that, while a particular brain region is sensitive to several oscillatory drivers, it will display a maximum sensitivity to a particular stimulation frequency (optimum frequency), and decreased sensitivities to drivers with frequencies outside of this range (e.g., [41–43]).

The NMM column consisted of two neural populations, connected in parallel (Figure 1). This combination allows the generation of a broadband EEG spectrum, from theta to gamma oscillations [35]. Each cortical population (resonant circuits) were defined to display different kinetics, such that they generated alpha (*α*) or gamma (*γ*) oscillations.

**Figure 1:**
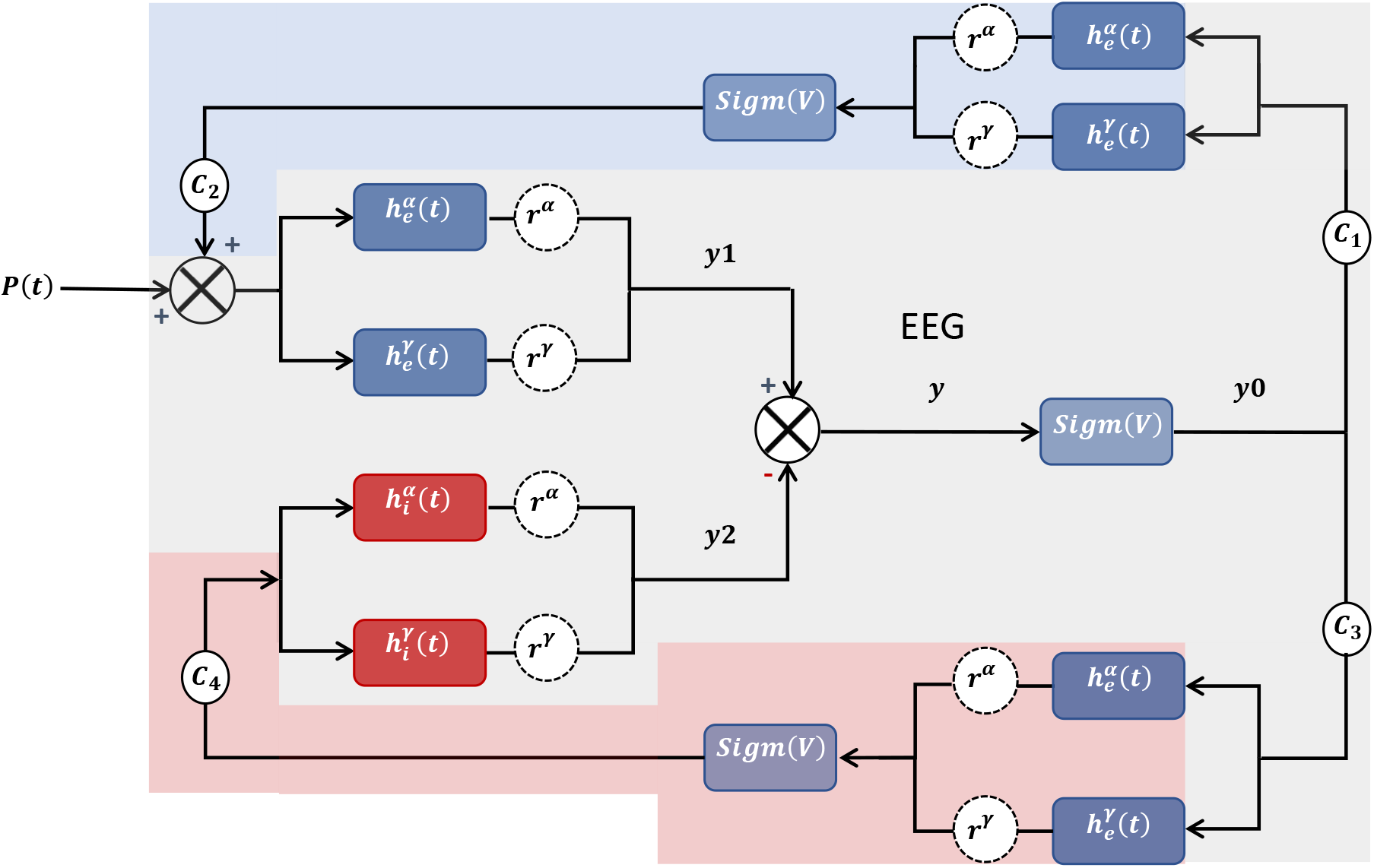
The modified Jansen-Rit NMM.The cortical column is made of two populations (*α* and *γ* neurons). Each population comprises three sub-population (types) of neurons; excitatory pyramidal cells (gray shadow), excitatory inter-neurons (blue shadow) and inhibitory interneurons (red shadow). The parameters *r^α^* and *r^γ^* tune the contribution of each sub-population to the column such that when the column is pure *α r^α^* = 1 and when the column generates pure *γ* oscillations *r^γ^* = 1 and *r^α^* = 1. The postsynaptic (PSP) functions are split into 2 parts [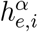 and 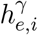] representing populations of *α* and *γ* kinetics, respectively.

The model includes excitatory/inhibitory interneurons interacting with pyramidal cells and uses two non-linear transformations. The first represents the site of generation of action potentials, and is a wave-to-pulse conversion. This function relates the mean firing rate of neurons to average post-synaptic depolarization. The other transformation is a pulse-to-wave conversion implemented at a synaptic level, which models the average post-synaptic response as a linear convolution of the incoming spike rate [35].

Each neural population comprised three sub-population (types) of neurons; excitatory pyramidal cells, excitatory inter-neurons (spiny stellate cells) and inhibitory interneurons (Figure 1). In this circuit, excitatory output of the pyramidal cells is sent to both types of interneuron sub-populations, which provide feedback loops to the pyramidal cells. The extrinsic input *P*(*t*), which is composed of noise plus any stimulus-driven input is received by the pyramidal cells. The extrinsic input of the column represents a pulse density with arbitrary units [40].

Since each population (*α* and *γ*) have both excitatory and inhibitory sub-populations, four postsynaptic generating functions are obtained for the column: 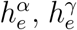 and 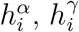 (Table 1).

**Table 1:** Postsynaptic generating functions for each neural population (*α*, and *γ*), and their corresponding excitatory and inhibitory sub-populations

| Type of sub-population | $\alpha$ population | $\gamma$ population |
| --- | --- | --- |
| Excitatory | $h_e^\alpha(t) = A^\alpha a^\alpha t e^{-a^\alpha t}$ | $h_e^\gamma(t) = A^\gamma a^\gamma t e^{-a^\gamma t}$ |
| Inhibitory | $h_i^\alpha(t) = B^\alpha b^\alpha t e^{-b^\alpha t}$ | $h_i^\gamma(t) = B^\gamma b^\gamma t e^{-b^\gamma t}$ |

$A^{\alpha,\gamma}$ , $B^{\alpha,\gamma}$ represent the maximum amplitude of the excitatory and inhibitory PSP for each sub-population and $a^{\alpha,\gamma}$ , $b^{\alpha,\gamma}$ are the so-called lumped parameters that take into account all the distributed delays in the dendritic network [36].

The parameters *a^α^, A^α^, b^α^, B^α^* were obtained from anatomical data [38,40], and were determined such that the resonance frequency of the neural populations were in the *α* frequency range of the EEG (≈10 Hz), to represent the visual cortex. The parameters (*α^γ^, A^γ^, b^γ^, B^γ^*) are those described in [35] to represent local neuronal peacemakers [44]. Both sets of parameters used for the PSP generation of the two populations are presented in Supplementary Tables 1 and 2.

The relative proportion of each population within the cortical column is controlled by the “ratio” parameter *r*; 0 < *r* < 1 (Figure 1) [35]. As a consequence, when *r* = 1, the NMM will generate *α* oscillations and when *r* = 0 oscillations will be in the *γ* range. We define *r^α^* as the proportion of the *α* generating population, noting that *r^γ^* = 1 − *r^α^*.

In the model, the EEG-like activity is represented by *y* (Figure 1), which is computed as the linear sum of the outputs from each neural population of the column; where *y*1^*α*^ and *y*1^*γ*^ are the outputs of the of the excitatory inter-neurons for both *α* and *γ* populations respectively, and *y*2^*α*^, *y*2^*γ*^ are the outputs of the of the inhibitory inter-neurons for both *α* and *γ* populations respectively. We can consider this EEG-like activity to represent the activity of the neural generators of the EEG, without taking into account the volume conduction or the head geometry. Moreover, the output (mean firing rate) of the pyramidal cells is represented by y0 (Figure 1).

The implicit assumption for the two-populations model is that every population expresses the same cyto-architectonic structure, having on average the same inputs and using identical constants *C_k_* with *k* = 1, 2, 3, 4 as in [35]. The intra sub-population connectivity constants *C*_1_, *C*_2_, *C*_3_ and *C*_4_ were determined using anatomical information [38] and represent the average of the number of synaptic connections between sub-populations (Figure 1). All the connectivity constants are expressed as a fraction of *C* (Supplementary Materials Table 3).

*Sigm*(*V*) denotes a nonlinear sigmoid function which transforms the average membrane potential of a neural population into an average firing rate 1.The function Sigm remains the same for each sub-population, for both excitatory and inhibitory branches of the cortical column. The Sigm function (equation 1) shapes classical properties of neurons: thresholds and saturation for the generation of action potentials [40].

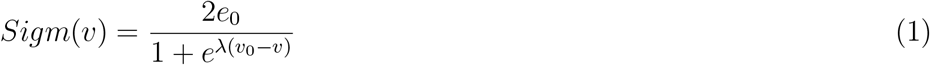

In Equation 1, *e*_0_ determines the maximal firing rate of the population, λ controls the steepness of the sigmoidal function, and *v*_0_ is the post-synaptic potential corresponding to the 50% of the firing rate (*e*_0_). Moreover, *v*_0_ can be either viewed as a firing threshold or as the excitability of the populations, whereas *v* is the average pre-synaptic membrane potential. The values of these parameters were empirically determined in [45].

#### 2.1.2. Coupling two modified Jansen-Rit NMM columns

Inter-column connectivity is defined as in [35] (Figure 2). Both columns of the model received uncorrelated extrinsic noise inputs (*n*1(*t*) and *n*2(*t*)) representing the background noise in the cortex. Noise was sampled from a Gaussian distribution with a mean of 221 and standard deviation of 31 [38, 40].

**Figure 2:**
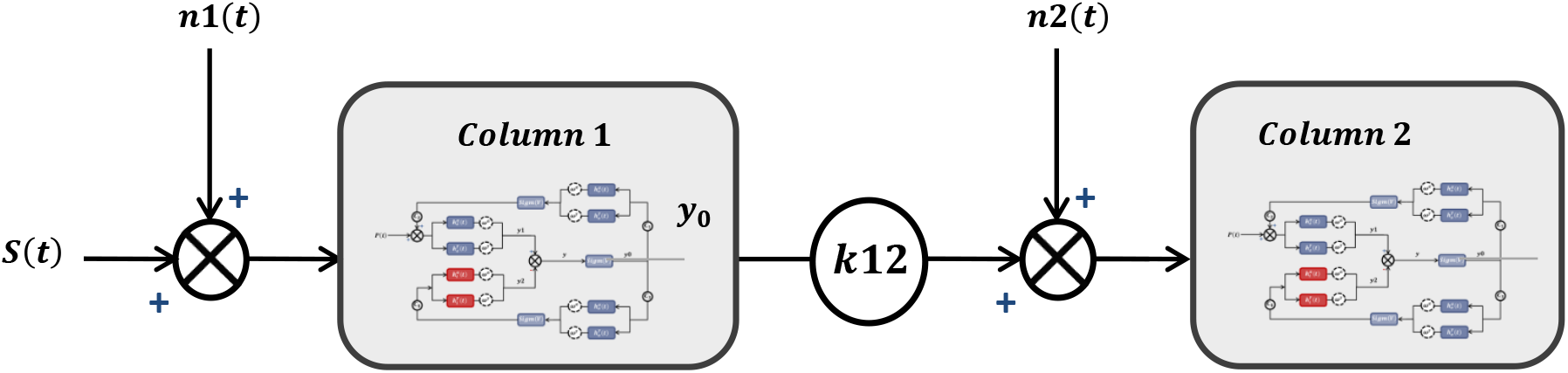
Schematic representation of the Jansen-Rit model consisting in two cortical columns of two populations each. Areas are coupled in cascade (unidirectionally from column 1 to column 2). Noise inputs *n*1(*t*) and *n*2(*t*) targets columns 1 and 2 at the same time, respectively, but only the first column was targeted by the sinusoidal driven input *S*(*t*). The coupling strength *k*12 mediates the connectivity between columns (equation 2).

Only the first column received a sinusoidal input *S*(*t*), which was added to the noise, *n*1(*t*). The decision to use a sinusoidal function as input to the model, simulates the effect of sinusoidally-varying stimulation of EEG oscillations, which include the effect of the terminating phase of a driving stimulus (phase of the stimulation at the offset) on the persistence of the entrainment [27].

The coupling strength from column 1 to column 2 was controlled by the coupling coefficient [38], *k*12, which attenuates the output of the first column (*y*0) before feeding into the second column (Figure 2). *k*12 was initially set as a value between 0 and 1, and was recalculated in every iteration [35] (noted as *k*12*(*n*)) to adjust for the total variance of the system as shown in Equation 2:

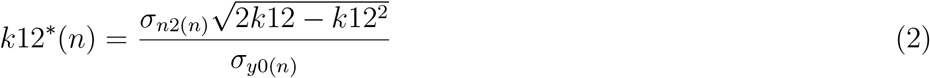

where *σ*_*n*2(*n*)_ is the variance of the noise input to column 2 and *σ*_*y*0(*n*)_ is the variance of the output from column 1, both in the nth iteration. This allowed us to specify a coupling coefficient *k*12*(*n*) bounded between 0 (no coupling) and 1 (no extrinsic input noise). The system of equations governing a 2-population 2-column model are given in Supplementary Information.

All the simulations were implemented in Matlab (2014b). The stochastic Euler expansion method was used to solve the system of ordinary differential equations governing the behaviour of the units, with initial conditions set to zero. Parameter values can be found in Supplementary Information 1, 2, 3 and 4. The system was approximated via an Euler expansion with an integration step of Δ*t* =1*ms* [46] as it has been shown that the numerical solution generated with this method has a distribution that resembles the distribution of the exact solution in the mean square sense (mean-square error of order Δ*t*^2^) [47]. In a pilot study, we found that doubling or halving the length of the integration step did not effect the results.

### 2.2. Simulations

Initially, we tested the effects of entrainment on a single column (equivalent to have coupled columns with *k*12 = 0).

- The power of alpha oscillations of column 1 as a function of *α* proportion *r^α^*, when a combination of white noise and an 11 Hz oscillation was the input of column 1 (section 3.1).
- The power spectrum of the oscillatory activity of a cortical column as a function of *r^α^* and the frequency of the driving force (section 3.1).
- The duration of the entrainment persistence as a function of *r^α^* (section 3.2). The methodology followed to compute the persistence duration is described in Methods section 2.3. Next, we tested propagation of the entrained signal between 2 columns.
- The power of the entrainment of column 2, as a function of *r^α^* of the column 2, the *r^α^* of column 1, and *k*12, when the first unit received an 11-Hz oscillatory input (sections 3.3 and 3.4).
- The duration of the entrainment persistence of column 2, as a function of *r^α^* of the column 2, the *r^α^* of column 1, and *k*12, when the first unit received an 11 Hz oscillatory input (sections 3.5).
- The effect of the terminating phase of the sinusoidally input of column 1 on the persistence duration of column 2, when *r^α^*=0.9 for both columns, and *k*12=0.9 (section 3.6).

Unless explicitly stated otherwise, simulations were run for a total of 5000 time steps (5 s). The stimulus lasted for 2745 ms, such that the terminating phase of the oscillation was *π*. The model output (*y*) was divided into three stages: baseline pre-stimulus (the 1-second period just preceding the stimulus onset), SSVEP interval (defined as the time interval between 0.5 s after the stimulus onset and the stimulus offset), and post-stimulus stage (from the stimulus offset to the end of the simulation.

Simulations were performed such that *r^α^* was varied between 0.1 and 0.9, in steps of 0.1. We simulated 1000 trials for each *r^α^* value. From the pool of traces obtained for each *r^α^*, 50 trials were randomly sampled without replacement. This operation was repeated 30 times to simulate 30 individuals with 50 trials each. The power spectrum of the ongoing oscillations (pre-stimulus interval, 1 s before the stimulus onset), and the neural entrainment (time interval between 0.5 s after the stimulus onset and the stimulus offset) were computed using the discrete Fourier transform. The mean power spectrum [48] of both baseline and entrainment was computed for each *r^α^* in each subject (50 trials). Following, the corresponding sample mean (30 subjects) was obtained.

### 2.3. Persistence of the entrainment duration computation

We computed the persistence duration of the model output after the stimulus offset as described in [27] (Figure 3). In summary, for each experimental condition (phase termination), 50 trials were randomly sampled without replacement 30 times from the entire data, such that the response of 30 subjects with 50 trials each was simulated. Trials belonging to the same subject were averaged in the time-domain (simulating an experimental condition in a particular individual) (Figure 3 A). The averaged signals were narrow-band filtered (±1 Hz around the stimulation frequency), using a zero-phase shift Butterworth filter of order 8 (time constant of 0.018 s for a 10 Hz oscillation). As described by Otero et al. [27], this procedure did not affect the evident oscillatory activity which succeeded the stimulus offset. Narrow-band filtering of the signal was performed to allow estimation of the physiological parameters of the entrainment (amplitude of the oscillations, duration of the persistence, among others) from the envelope of the time-domain averaged signal, which in turn was computed using the Hilbert transform. Computation of envelopes using the Hilbert transform is particularly recommended on narrow-band filtered signals [49–52].

**Figure 3:**
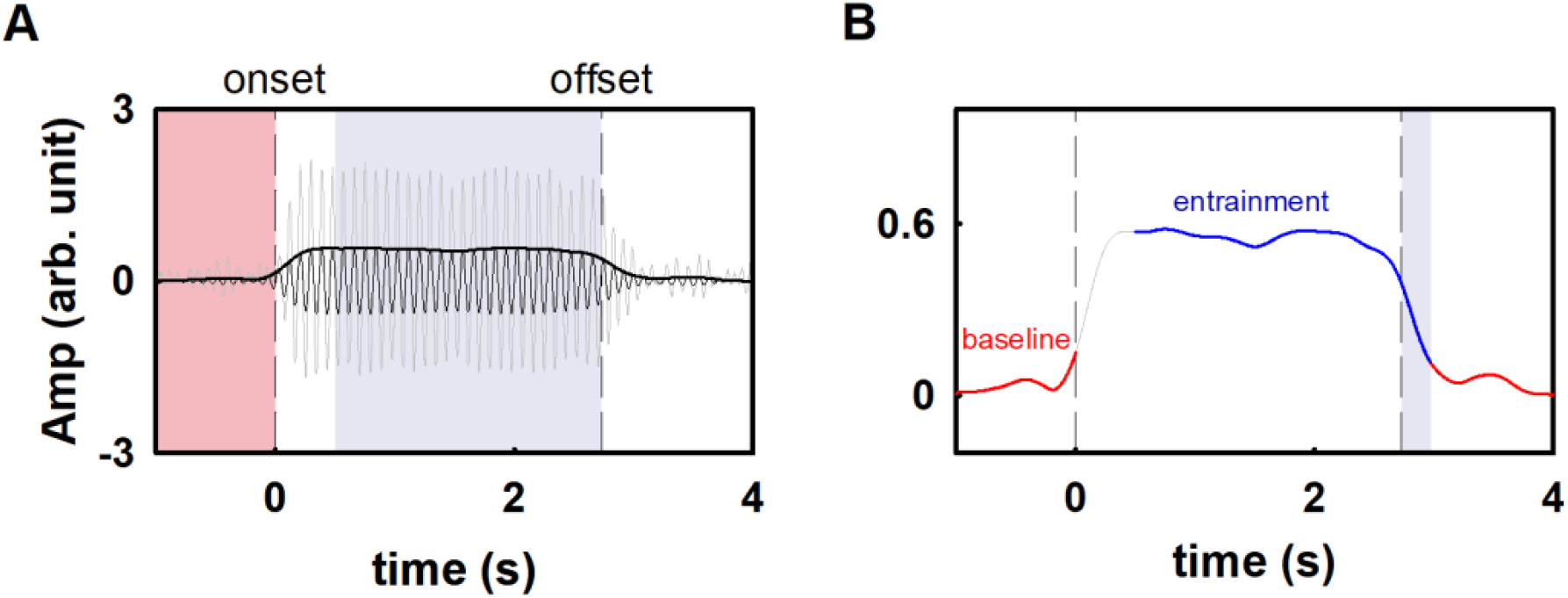
Summary of the pipeline for calculating the persistence duration (PD). **(A)** SSVEP signals are bandpass filtered ±1 Hz around the frequency of stimulation. The envelope of the SSVEP signal is extracted using the Hilbert transform, which results in an instantaneous estimate of the amplitude. Dashed vertical lines represent the stimulus onset and offset, respectively. **(B)** Time points beyond stimulus offset are classified into entrained (blue) or not entrained (red) regions. Persistence duration is defined as the time from stimulus offset where the signal is continuously classified as belonging to the entrained distribution.

We use the Hilbert function provided in Matlab to obtain Discrete-time analytic signal and its envelope. The time resolution of the Hilbert transform was calculated such that the Full-Width at Half-Maximum (FWHM) of the impulse response of the Hilbert transform corresponded to 7.8 ms.

Simulated signals were divided in (pre-stimulus interval, 1 s before the stimulus onset), and the neural entrainment (time interval between 0.5 s after the stimulus onset and the stimulus offset) (Figure 3 A). Individual classifiers were constructed based on the amplitude distribution of pre-stimulus (stimulus off, not entrained) and during 1 s of stimulation (SSVEP, computed in the time period between 1 and 2 seconds after the stimulus onset). This step allowed us to identify the boundary between the persistence of entrainment and baseline. This methodology followed a signal detection approach [53], where two distributions are discriminated: the “noise” distribution given by the baseline (in this case, the simulated ongoing EEG) and the “signal” distribution given by the entrainment (simulated SSVEP signal). Gaussian functions were fit to the instantaneous amplitudes of pre-stimulus and SSVEP, and the constructed normal distributions were used to calculate the probability of the post-stimulus amplitudes belonging to either baseline or entrained stages. The cutoff value for classification was the intersection between the baseline and the entrained distributions. Therefore, the boundary between the persistence of entrainment and post-stimulus baseline (cutoff value) was defined using a neutral decision criterion, where neither stage was favored [27]. The duration of the persistence of entrainment was defined as the time interval after the stimulus offset which was classified as belonging to the entrained stage, or conversely, that differed from baseline (Figure 3B).

## 3. Results

### 3.1. Simulation 1: Neural entrainment as a function of resonance frequency of the NMM

Initially, we investigate the effects of entrainment and persistence on a single-column model. In this section, the NMM column was tuned to exhibit different intrinsic oscillatory activity which was achieved by varying the relative proportion of *α* and *γ* populations. We observe the effect of an entraining frequency in the *α* range (11 Hz) on the columns dynamics. A detailed analysis of the intrinsic oscillations can be found in Supplementary Information.

In Figure 4, we illustrated examples of the activity generated by single columns with intrinsic oscillations in the gamma range (*r^α^* = 0.2, and *r^α^* = 0.3), as well as alpha oscillations (*r^α^* = 0.8, and *r^α^* = 0.9) when an 11 Hz driving force was applied. As expected, intrinsic oscillations were evident before the presentation of the stimulus. The frequency of these intrinsic oscillations depended on *r^α^*. Columns with low *r^α^* exhibited basal oscillations in the gamma frequency-range, columns with high *r^α^*) exhibited intrinsic oscillations in the alpha frequency range (Figure 4, middle panels). After the stimulus onset (time 0 s), amplitude of the intrinsic oscillation increased as compared to the pre-stimulus interval. The amplitude of the oscillation remained stable until the end of the stimulation. In other words, the 11 Hz driver entrained the intrinsic oscillation of the NMM column (Figure 4, left panels). Consequently, power spectra had a clear peak at the frequency bin corresponding with that of the oscillatory input (Figure 4, right panels). Hereafter, this spectral peak will be referred as the alpha power of entrainment.

**Figure 4:**
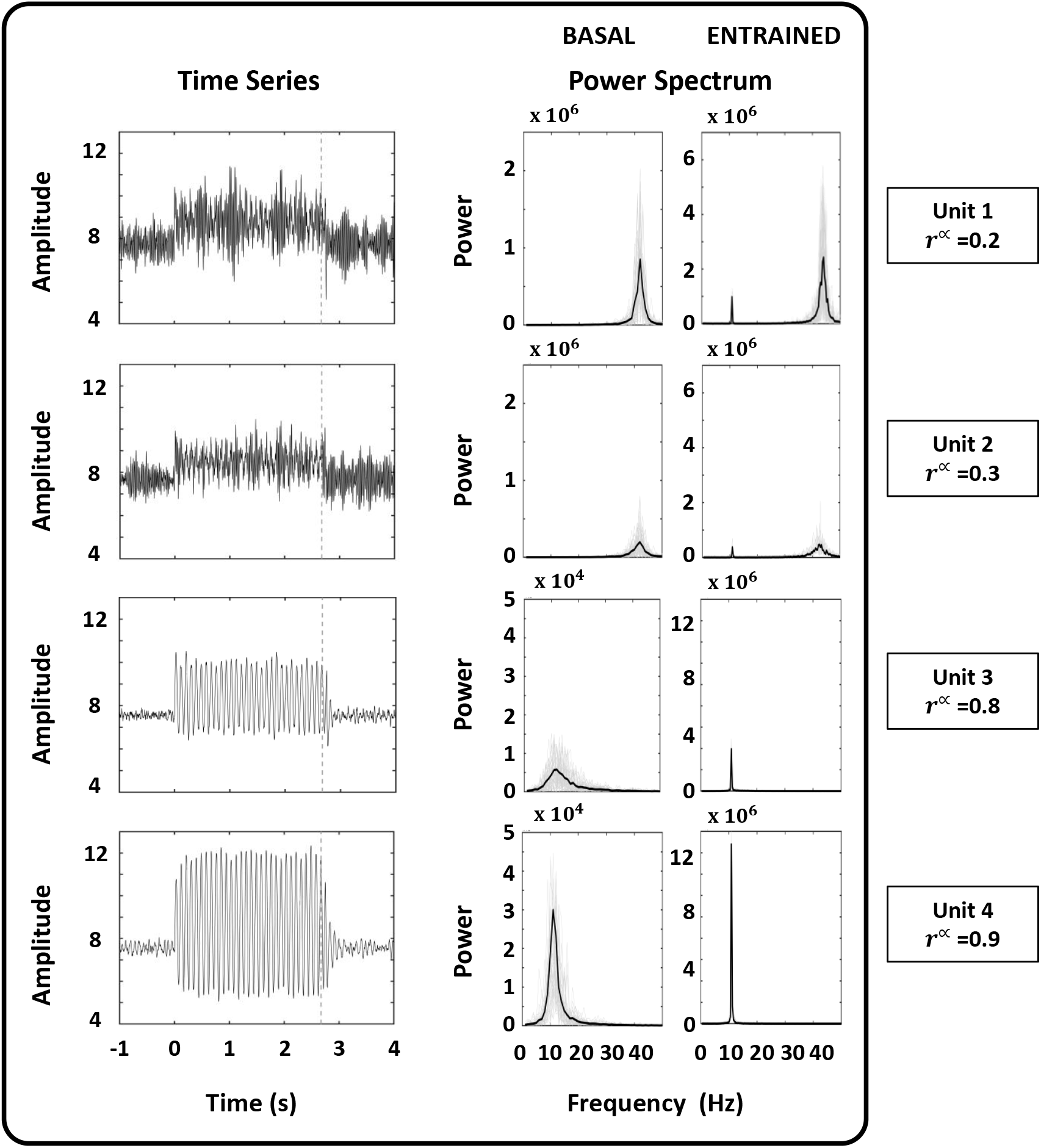
Examples of the response of single NMM column to oscillatory external drivers of 11 Hz. The examples correspond to *α* proportion values of *r^α^* = 0.2 and *r^α^* = 0.3, which resulted in the generation of gamma oscillations, and *r^α^* = 0.8 and *r^α^* = 0.9, which resulted in alpha oscillations. For each *r^α^*, the time series and power spectrum of the column activity are presented. The time series represent the time-average of all the trials. The onset of the stimulation corresponds to time zero, whereas the stimulus offset is represented by a dashed vertical line. Power spectrum was computed for both basal and entrained conditions.

Alpha power at the entraining frequency was found to be directly related to *r^α^*, such that higher values for *r^α^* were associated with greater power at the entraining frequency. Based on the results presented above, the parameter *r^α^* should be interpreted as the ability of the columns to be entrained to extrinsic alpha oscillations.

A comprehensive representation of the effect of *r^α^* on the power of the alpha entrainment elicited by a 11 Hz driving force is illustrated in Figure 5. When the proportion of neurons tuned to alpha oscillations was less than 0.5, the power of alpha oscillations were relatively small. However, for *r^α^* between 0.1 and 0.5, the mean alpha power was always higher than zero. For *r^α^* higher than 0.5, the alpha power systematically increased, although a robust entrainment was only obtained when *r^α^* was 0.9.

**Figure 5:**
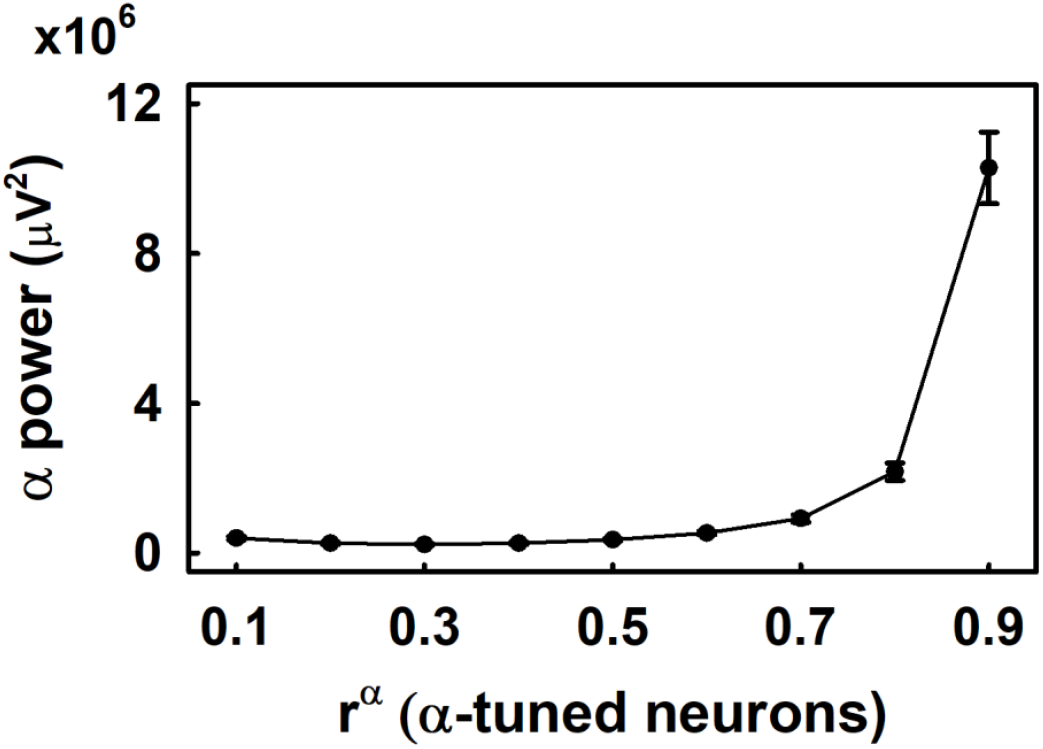
Alpha power as a function of the *α* proportion (*r^α^*) in a single-column NMM. The frequency of the driver was 11 Hz and *α* proportion *r^α^* was varied from 0.1 to 0.9 in steps of 0.1. Values represent the mean ± standard deviation of power, calculated from 50 simulations of the neural activity.

The results presented above were expanded to analyze the combined effect of *α* proportion (*r^α^*) and the frequency of the driving force on the power spectrum of the oscillatory activity of a NMM. To this end, the frequency of the oscillatory input (S(*t*), Figure 2) was varied from 1 up to 60 Hz, in steps of 1 Hz. Simultaneously, the *α* proportion was varied between *r^α^* = 0.1 and *r^α^* = 0.9, in steps of 0.01.

The most prominent feature of the model dynamics was the presence of two clear spectral peaks. They were obtained when the frequency of the driving force corresponded to the resonance frequency of the neural populations comprising the column, i.e., 11 and 43 Hz, respectively (Figure 6). This way, the maximum alpha power was obtained when *r^α^* was between 0.7 and 0.9 and the column was stimulated with the 11 Hz driver. Likewise, the maximum gamma power was obtained when *r^α^* was between 0.1 and 0.3 and the column was stimulated with the 43 Hz driver. The power of both alpha and gamma oscillations systematically decreased as the frequency of the driver moved away from the respective resonance frequency.

**Figure 6:**
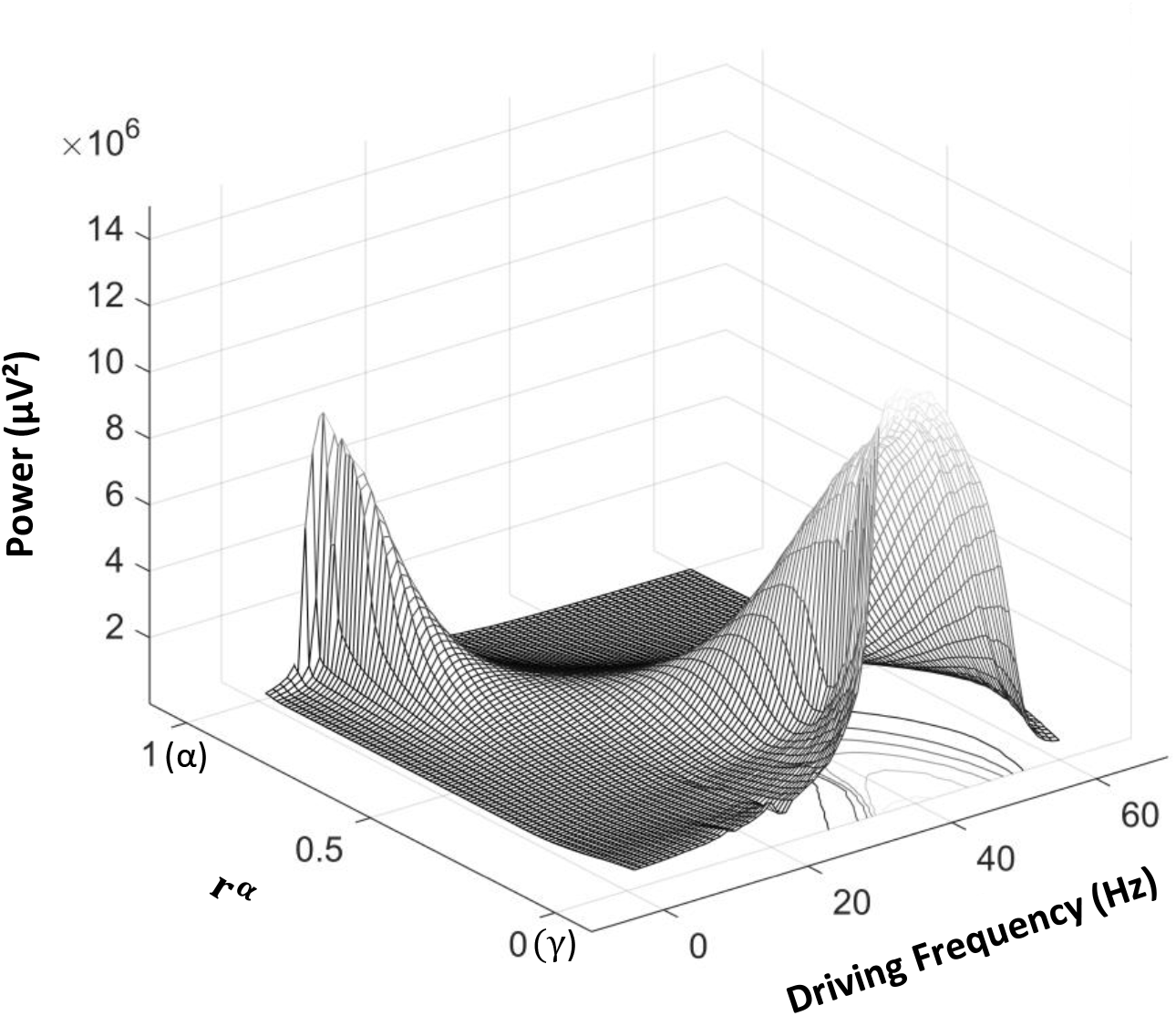
Resonance phenomenon. Entrainment of a single-column NMM as a function of the frequency of the driving force and the proportion of neurons tuned to alpha oscillations (*r^α^*). Alpha proportion *r^α^* was varied from 0.1 (*γ*) to 0.9 (*α*), while the driving frequency was varied from 1 to 60 Hz. Although the entrainment could be generated at any frequency, maximum responses were only obtained at those driving frequencies coinciding with the alpha or gamma band resonances.

We found that the maximum power spectrum of gamma entrained oscillations was higher than that of the alpha oscillations. This was not a consequence of a difference in the relative number of neurons responding to the stimulation, since the relative proportion of gamma-tuned neurons for *r^α^*=0.1 was equal to that of *α*-tuned neurons for *r^α^*=0.9. The differences in the power of alpha and gamma entrainment results from the differences in the post-synaptic (PSP) functions of the excitatory and inhibitory neurons comprising the *α* and *γ* populations, respectively (see Supplementary Information for details), where PSP amplitudes are greater for the *γ* population than for the *α* population.

### 3.2. Simulation 2: Persistence of the entrainment as a function of the resonance frequency of the NMM

The behavior of the entrainment after the stimulus offset, when the NMM column was tuned to *α* oscillations (Figure 4), resembled those described in previous studies [27, 29, 32, 33]. That is, the amplitude of the alpha oscillation did not return to baseline immediately after the stimulus offset but persisted for several hundred milliseconds.

For *r^α^* between 0.1 and 0.7, the alpha oscillatory activity of the NMM outlasted the stimulus offset for less than 100 ms, i.e., in less of a cycle of the sinusoidal input. Nevertheless, the mean PD was 270 ms when the *r^α^* of the NMM was increased to 0.9 (7). This PD was approximately three cycles of a typical alpha oscillation, and is comparable to those obtained in electrophysiological experiments [27–33].

### 3.3. Simulation 3: Propagation of the entrainment between coupled columns

Next we added a second column, coupled as described in Methods (section 2.1.2), to investigate the effects of resonance on the propagation of an entraining signal. Examples of the time evolution and spectral features of the oscillatory activity of identical columns coupled in cascade (with unidirectional coupling from column 1 to column 2) are presented in Figure 8. As expected, a clear entrainment of column 1 was obtained, when *r^α^* = 0.9 and a 11 Hz driving force was presented, (see Figure 8 A). The inter-column coupling coefficient (*k*12) was set to 0.5 (equation 2) and both basal oscillations and entrainment propagated. While the power of the basal alpha-oscillation of column 2 was increased as compared with that of column 1, the entrainment of column 2 was reduced in comparison with that of the first column.

A different behavior was obtained when columns were tuned to gamma oscillations (*r^α^* = 0.2) and the first columns received an 11 Hz driving force (Figure 8 B). In this condition, the alpha entrainment in the first column was not optimal (a substantial increase of the alpha band power was not obtained). Furthermore, the power of the gamma oscillations of the first column during the stimulation interval was similar to that of the basal stage (Figure 8 B).

### 3.4. Simulation 4: Effect of the coupling strength in the propagation of the entrainment

We investigated the effect of the *α* proportion on the alpha power of the second column of the pair, as a function of *r^α^* of column 2, and *k*12. The first column was tuned to alpha (*r^α^* = 0.9) or gamma oscillations (*r^α^* = 0.2), and received an 11 Hz oscillation input. As mentioned above, both cortical columns received extrinsic noise inputs.

The scenario in which columns 1 is tuned to alpha oscillations (*r^α^* = 0.9), and therefore is entrained by the driving force, is presented in Figure 9 (left panel). In this situation, when *r^α^* of column 2 varied between 0.1 and 0.7, the alpha power of the entrainment remained relatively constant and did not depended on the coupling strength *k*12 (Figure 9, left panel). Higher alpha power were consistently obtained as *r^α^* increased from 0.7 up to 0.9. Steeper increases in alpha power were obtained as *k*12 increased. Furthermore, the effect of *k*12 was not linear.

In the case that column 1 was tuned to gamma band oscillations *r^α^* = 0.2 (Figure 9, right panel), the alpha power of the second column did not varied as a function of *α* proportion in the entire range of *r^α^* we tested. This behavior was observed for any of the three *k*12 tested values.

### 3.5. Simulation 5: Propagation of the persistence of the entrainment

The PD of the entrainment of column 2 was analyzed as a function of the *r^α^* of the first column, *r^α^* of the second column, and *k*12. The results are illustrated in Figure 10.

We performed a two-way factorial ANOVA to analyze the effect of *r^α^* of column 2, and *k*12 on the PD. The factor *r^α^* of column 2 had nine levels (from 0.1 up to 0.9, in steps of 0.1), whereas factor *k*12 had three levels (0.1, 0.5, and 0.9). Both *r^α^* of column 2, and *k*12 had statistically significant effect on the duration of the persistence (F=127.72, p<0.05; and F=41.04, p<0.05 for the effects of *r^α^* and *k*12, respectively). Furthermore, the interaction *r^α^* and *k*12 also had significantly statistically effect on the duration of the persistence (F=4.65, p<0.05).

The PD of the entrainment of column 2, computed for a particular *k*12, did not vary when *r^α^* increased from 0.1 up to 0.4. For additional systematic increases in *r^α^*, longer PD were obtained (Fisher LSD post-hoc test, p<0.05). The PD of column 2, when the *r^α^* of this column increased from 0.1 up to 0.6, did not varied as a function of *k*12. For *r^α^* between 0.7 and 0.9, the PD increased as *k*12 varied from 0.1 to 0.5, and remained constant when *k*12 increased from 0.5 up to 0.9 (Figure 10).

In the case that the column receiving the 11 Hz was tuned to gamma oscillations (*r^α^* = 0.2), the coupling strength did not have any evident effect on the PD of the second column (Figure 10 right panel).

### 3.6. Simulation 6: Effect of the phase at the stimulus offset in the persistence duration

The persistence of the entrainment has been described experimentally as a function of the phase of the EEG at which stimuli were removed [27]. In this section, we will analyze if a modified Jansen’s model replicates this experimental results. We note that there are slight differences in the way the phase was controlled between simulated data in this research and EEG recordings in [27]. In the experimental study, the phase angles of the EEG at the time the stimulus ended were determined retrospectively (a posteriori). This implied that a particular experimental condition (phase of the stimulus at the offset) did not correspond to a particular phase angle of the EEG oscillation but were uniformly distributed in a range of phase angles (Figure 5 in [27]). By contrast, phase angles were determined beforehand (a priori) in the simulated data, hence the sinusoidal inputs corresponding to an specific phase condition finished at exactly the same phase angle.

We restricted the analysis to scenario in which the optimal entrainment persistence was obtained (*r^α^* = 0.9 for both columns, and *k*12=0.9) (Figure 10). Unlike previous simulations, the 11 Hz sinusoidal inputs ended at different phases. In other words, inputs corresponding to different terminating phases had different duration. The shortest stimulation was 2700 ms, which corresponds with 30 cycles of a sinusoidal function of frequency 11 Hz. Eighteen equidistant phases covering a cycle of the oscillation were sampled. In an 11 Hz sinusoid, this sampling is equivalent to have eighteen stimuli with different duration, in which the duration of stimuli representing two consecutive terminating phases varied in 5 ms. The PD corresponding to each terminating phase of the driving stimulus was calculated as described in section 2.3.

The PD of the alpha entrainment was highly dependent on the phase of the model output at the point the 11 Hz sinusoid was removed (Figure 11 A). The longest PD was found when the phase of the model output at the stimulus offset was *π*. The shortest PD were encountered in the opposite phase (phase 0). Therefore, while a group of terminating phases facilitated the persistence of the entrainment, others impede it.

To draw the analogy with the experimental results presented in [27] using EEG recordings, PD were pooled in two groups: persistence elicited by rising phases of the oscillation at the stimulus offset (phases ranging from 0 to *π*, and from 3*π*/2 to 2*π*) and persistence elicited by falling phases (phases range (*π*/2 to 3*π*/2). A t-test (p<0.05) was implemented to compared the mean PD obtained by the different group of phases (Figure 11 B). As a result, the persistence obtained when the phase of the EEG-like oscillation was towards the trough of the cycle (falling phases) was statistically significant longer than that obtained when the phase of the EEG-like oscillation was towards the peak of the cycle (falling phases) (t=14.51; p<0.05). This result is consistent with the experimental findings described by [27], although the specific phases that facilitate or impede entrainment were different between the model and the experiments. Possible reasons for this discrepancy will be addressed in Discussion section. 4.3.

## 4. Discussion

### 4.1. Summary

In this study, we followed David and Friston [35] in adapting the Jansen-Rit NMM [38,40] to construct a neural mass model capable of generating the entire EEG spectrum. This comprised a hybrid column with a mix of gamma and alpha tuned populations. Without loss of generality we investigated the effect of driving this column with external alpha oscillations on the entrainment of the NMM as a function of the alpha/gamma ratio of the column. Furthermore, we analyzed the propagation of the entrainment from one column to another, and the factors that allowed the entrainment to persist after termination of the oscillatory input. Our results indicate that the relative proportion of neurons of the NMM tuned to the frequency of the oscillatory driving force directly determines the strength of the entrained signal, and the duration of the persistence of the entrainment after the termination of the periodic input. When two NMM were coupled, the relative proportion of neurons tuned to the frequency of the oscillatory driving force in each NMM, as well as the coupling strength between columns were the factors that determined the propagation of the entrainment from one column to the other. Additionally, the persistence of the entrainment depended on the phase of the EEG-like oscillatory activity of the NMM at the time stimulus terminates.

Since a 11-Hz sinusoid (alpha band stimulation) was the driving force that served as input to the NMM, our study provides a plausible explanation for experimental results analyzing the entrainment of EEG alpha oscillation in response to periodic visual stimulation in the alpha band [14, 16, 25]. Remarkably, our results suggest that the internal functioning of the NMM, which in turn is defined by the cytoarchitecture of the cerebral cortex, is able to account for the persistence of the stimulus-driving alpha oscillation after the end of the sensory stimulation [24, 28–33], as well as the effect of the terminating phase of a sinusoidal light stimulus on the duration of the persistence [27]. Although we restricted the analysis of the entrainment to alpha oscillations, our results can be extrapolated to any particular frequency band of the EEG.

### 4.2. Frequency-dependent neural entrainment

The proportion of neurons tuned to a particular stimulus oscillatory frequency in a given brain region is difficult to be determined experimentally. Using simulations, the results presented in Figure 5 demonstrate that 10 percent of alpha tuned neurons in the NNM is sufficient to generate alpha entrainment. Nevertheless, the entrainment will be maximum only when the driver matches the resonance frequency of the oscillatory neural system (Figures 5 and 6). This is in accordance with previous studies [54,55], showing that the coding of stimulus periodicities in a particular brain area is performed by different neuronal populations, in which each neural ensemble is tuned to a given range of driving frequencies. The sensitivity of the area to oscillatory stimuli is then determined by the relative size of the different populations comprising the area, such that the resonance frequency of the cerebral region will correspond to the frequency at which the largest population forming the area is tuned to. This is evident in the oscillatory activity of cortical columns with different *r^α^* presented in Figure 5. In those columns, the frequency of intrinsic oscillations (the resonance frequency) is determined by the oscillatory properties of the neural population most represented in the cortical region. These results support the idea of that the amplitude of the stimulus-induced oscillation in a particular brain region, as a function of the frequency of the stimulation, is proportional to the size of the neural population tuned to the frequency of the sensory stimulus. [41,56].

Furthermore, the results presented in this study contribute to the understating of steady-state responses recorded from the scalp, i.e., EEG oscillations locked to the periodicity of the stimulation, which have relatively constant amplitude and phase over the stimulation interval. These oscillatory responses can be interpreted, at least partially, as the phase alignment of multiple EEG generators (multiple NMM) responding to the periodic stimulation [25]. In other words, while neural entrainment is the physiological process occurring at the level of EEG generators, the steady-state responses represent the observable manifestation of the entrainment. Consequently, the fact of having different number of neurons with a distinct resonance frequency is also a theoretical support for experimental data of scalp-recorded SSEP, which exhibits different amplitudes in response to sensory stimuli of equal modality and intensity, but presented at different frequencies [54, 55, 57]. This sensitivity to the stimulus periodicity is represented by the temporal modulation transfer function of the oscillatory response (the amplitude of the steady-state responses as a function of the frequency of the driving force eliciting the oscillation). This is a typical inverted V-shaped function, where the frequency at which the maximum amplitude is obtained (best modulation frequency of the steady-state response) can be assumed as the resonance frequency of the underlying neural generators. For frequencies above and below the best modulation frequency, the amplitude of the steady-state response is proportional to the size of the neural populations tagging the frequency of the sensory stimulation [41].

In fact, the size of the neural population responding to the sensory stimulation could determine the sensitivity of the neural network to the stimulus periodicity, i.e., the resonance frequency of the steady-state generator, and therefore the best modulation frequency of the scalp-recorded brain oscillation. This topic has been addressed in a recent study [58], which demonstrated that the driving frequency which elicits the maximum amplitude of steady-state responses in the visual cortex is inversely correlated with the area of the visual field covered by the stimulation. Assuming that expanded stimuli activate larger neural population, this result indicates that the resonance frequency of the oscillatory neural network is ultimately determined by the the size of the network [58]. This can be extrapolated to the resonance frequency of steady-state responses of different modalities. In the human brain, the volume of the primary visual, somatosensory and auditory primary cortices are 23 *cm*^3^, 13 *cm*^3^ and 3 *cm*^3^, respectively (e.g, [59–61]). Considering that the cytoarchitecture remain constant across these sensory cortices, the highest amplitude of the SSVEP, as compared to that of other sensory modalities, represents the largest size of the visual cortex relative to that of the other primary sensory regions.

Furthermore, our modeling of entrainment suggest that, in the case that different neuronal populations tagging different regions of the spectrum have the same size, the dynamic of the ionic current generating the PSP might be critical for the oscillatory activity of the neurons. This is evident in the power of the alpha and gamma entrainment of NMM columns with equivalent proportion of neurons tuned to the particular driving frequency (when *r^α^* was 0.9 and 0.1, respectively, Figure 6). This difference is a consequence of the greater post-synaptic (PSP) amplitude of the gamma-tuned neurons as compared to that of the alpha-tuned neurons, which in turn depend of the ratio between the amplitude of the inhibitory and excitatory PSP (Table 1).

### 4.3. Mechanism of the persistence of entrainment

Experimental studies in different sensory modalities have consistently demonstrated that the SSEP persist after the stimulus offset for times equivalent to three periods of the entraining stimulus, approximately [27–30]. Our modeling results shed light on the conditions needed for the entrainment to outlast the stimulus offset.

When the proportion of alpha-tuned neurons (*r^α^*) of the NMM is lower than 0.7, the entrainment elicited by an 11 Hz driving force outlast the stimulus offset in less of a cycle of the stimulus oscillation, i.e., the persistence duration is negligible (Figure 7). This indicates that the entrainment of a given NMM will persist after the stimulus offset only when more than 70 percent of its neurons are tuned to the input periodicities. To reproduce empirical observations of persistence, it was necessary that at least 90 percent of the neural population has resonance frequency equal to that of the driving force (Figure 7). In this regard, it is important to mention that the variations in the entrainment persistence as a function of *r^α^* (Figure 7) resemble the relationship between the power of entrainment and *r^α^* (Figure 5). This indicates that robust entrainment is needed for the oscillation be able to outlast the stimulus offset, which in turn is achieved when a sufficiently high amount of neurons of the brain region has resonance frequency equal to the frequency of the driving force. These results strongly support experimental results showing that driving forces with frequency outside the EEG alpha band elicit on-responses at the beginning, and off-responses at the end of stimulation instead of alpha entrainment [62].

**Figure 7:**
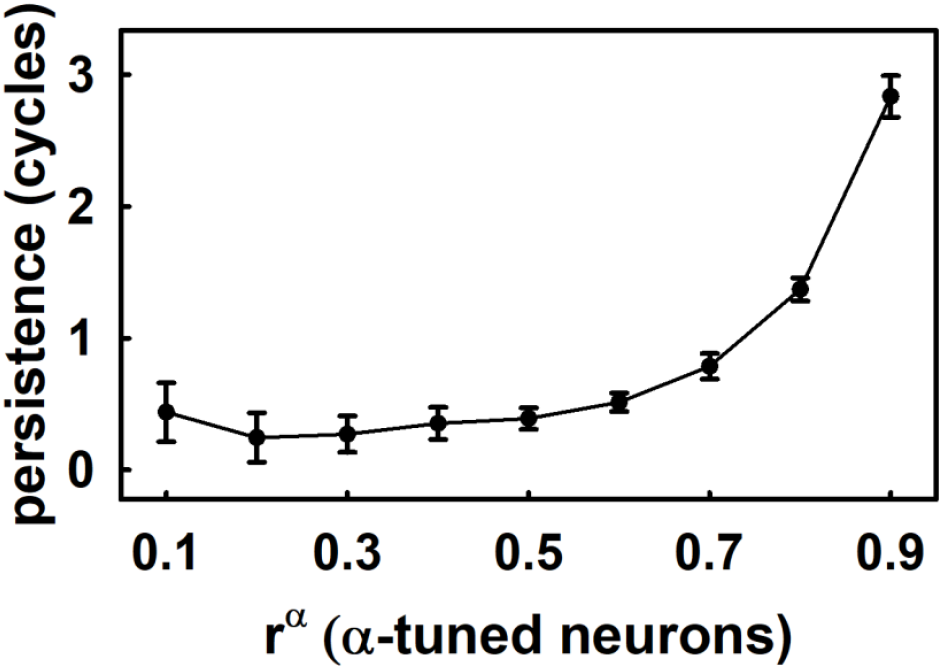
Effect of *r^α^* on the persistence duration of the entrainment elicited by a driving force of frequency 11 Hz. The plot represents the mean ± standard deviation of the persistence, calculated from 30 simulated subjects.

**Figure 8:**
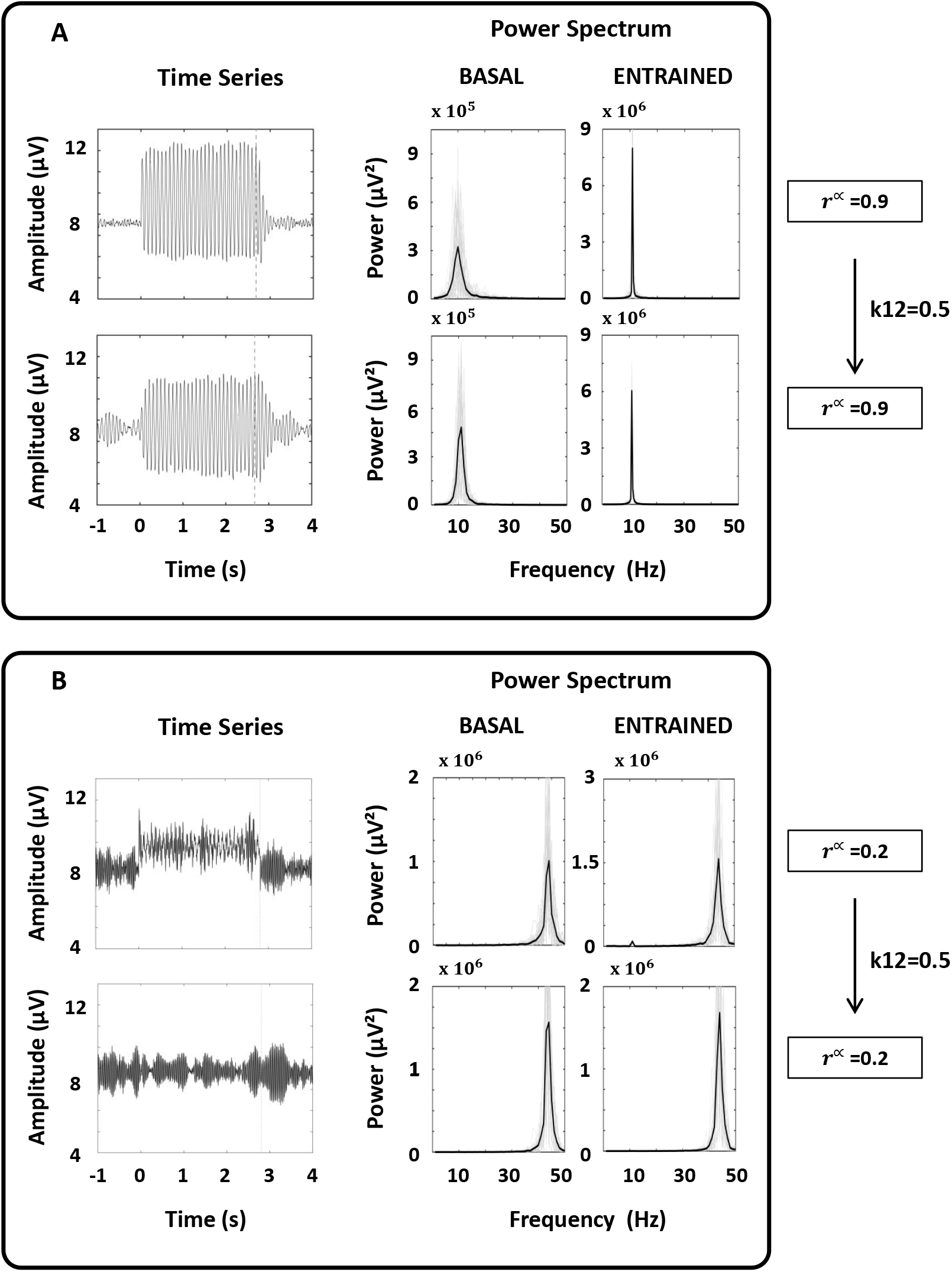
Examples of the entrainment of identical columns with unidirectional coupling from column 1 to column 2. The first column was targeted with a driver at 11 Hz. Coupling strength from column 1 to column 2 is *k*12 = 0.5. Time series and power spectrum is shown, in the basal (left) and the entrained (right) conditions. (**A**) Two identical columns with *α* proportion *r^α^* = 0.9. (**B**) Two identical columns with *α* proportion *r^α^* = 0.2.

**Figure 9:**
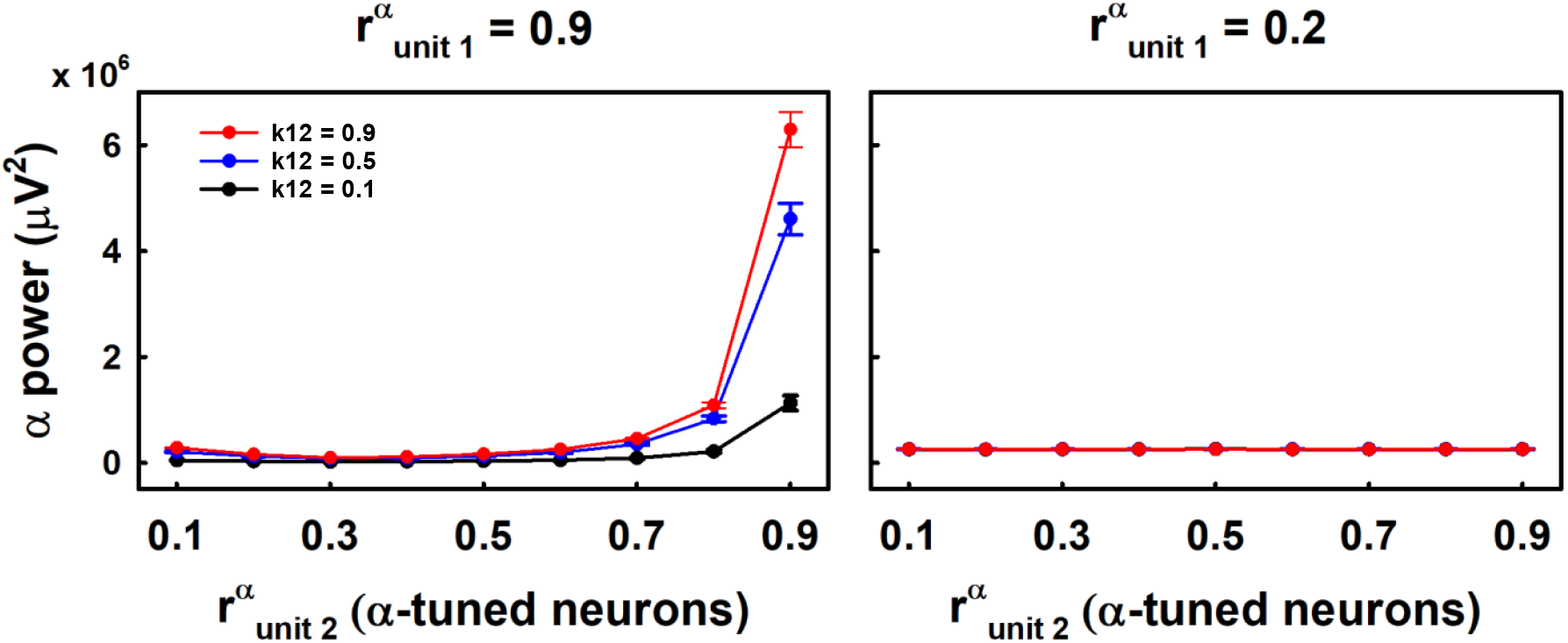
Effect of coupling strength *k*12 on the propagation of the entrainment as a function of the *α* proportion *r^α^* of the second column. Three values of *k*12 are presented: black lines (*k*12 = 0.1), blue lines (*k*12 = 0.5) and red lines (*k*12 = 0.9). The alpha power of a second column coupled to a first column receiving an oscillatory input at 11 Hz is shown. **(Left panel)** Alpha proportion of the first column was *r^α^* = 0.9. **(Right panel)** Alpha proportion of the first column was *r^α^* = 0.2.

The persistence of the entrainment can be explained by the presence of feedback loops (reverberant circuits) in the functioning of cortical columns. Considering the circuitry of a NMM when *r^α^*=0.9 (Figure 1), the output of the pyramidal cell *y*0 just after the oscillatory input terminates will be high enough to make the feed-back loops to have significant influences in the post-synaptic potential of the pyramidal cells. Consequently, both *y*1 and *y*2 (the output of the excitatory inter-neurons and the output of the inhibitory inter-neurons, respectively) will present values above those computed when the input of the model is exclusively composed by noise (baseline). The integration of these excitatory and inhibitory post-synaptic potentials (summation of *y*1 and *y*2) then results in a postsynaptic potential *y* higher than the noise floor. The entrainment of the NMM will persist as long as *y* remains above baseline. In the subsequent cycle of the loop, lower *y*1 and *y*2 are systematically obtained in absent of stimulation, such that *y* return to baseline at times which correspond with three periods of an oscillation with frequency equal to the resonance frequency of the alpha-tuned population.

### 4.4. Persistence duration of the entrainment depends on the phase of the EEG-like signal at the stimulus offset

The results replicated empirical findings which demonstrated that the phase of the stimulus offset determined the duration of persistence (Figure 11) [27]. Specifically, the rising phases of the stimulus offset (EEG phases towards the trough of the alpha-oscillation) showed to facilitate the persistence of entrainment while falling phases impede it, i.e., they induce longer and shorter persistence duration, respectively [27]. Additionally, the in silico model allowed us to expand the resolution of the phase information meaning that we could analyse the effect of stimulus offset during 18 phases (compared to 4 in the experimental study [27]).

Considering the different nature of the oscillatory activity elicited by the human brain and that of the NMM, direct comparison of the results is difficult. Nevertheless, this study, as well the results described in [27], provide evidence that the behavior of the entrainment after the stimulus offset was sensitive to the phase at the stimulus offset. These can be understood as the excitation/inhibition state of the neural network at the time the stimulus terminates, and may vary as a function of arousal and the level of engagement to a particular task [63]. These authors implemented a cortical oscillator network model which included a thalamo-cortical loop as the main circuit involved in the generation and maintenance of alpha oscillations, and simulated the oscillatory activity in three different states: eyes-open, eyes-closed, and task-engaged. They observed that stimulation enhanced endogenous oscillations both during and immediately after stimulation and that enhancement depended on the brain state, results that were corroborated by experiments of neuro-modulation with repetitive transcranial alternating current stimulation.

### 4.5. Propagation of the entrainment

The finding that entrainment is able to propagate between units in the model further corroborates experimental results [64] in which the presentation of simultaneous, amplitude-modulated audio-visual input in the theta band, enhanced the episodic (associative) memory for these audio-visual streams. The theta-specificity of these memory effects suggested that the sensory entrainment reaches downstream memory areas that are known to resonate at theta frequency, such as the hippocampus [14]. Hence, the entrainment by external rhythmic stimulation was found to propagate beyond the input area (supra-modality of entrainment) [64]. The double column NMM, in which units are connected in cascade (from cloumn 1 to column 2), can be understood as a representation of hierarchical organisation of neural sub-populations within a cortical region, or different areas.

When a driving force is presented to the first column of the pair of connected units, the propagation of the entrainment depended on three factors. First and foremost, the initial (sending) unit needs to be stimulated at its resonance frequency. Furthermore, it is necessary that the oscillatory activity of the first unit matches (or it is near to) the resonance frequency of the second unit. If these conditions are fulfilled, the propagation will depend on factors controlling the information transfer between units, which is represented in the model by *k*12 (Figure 2). In a physiological system, this parameter can be interpreted as the density of connections between the regions, in combination with the type and density of ion channels mediating the synaptic transmission. The rules governing the propagation of the steady region of the entrainment also regulate the propagation of the persistence (Figure 10).

**Figure 10:**
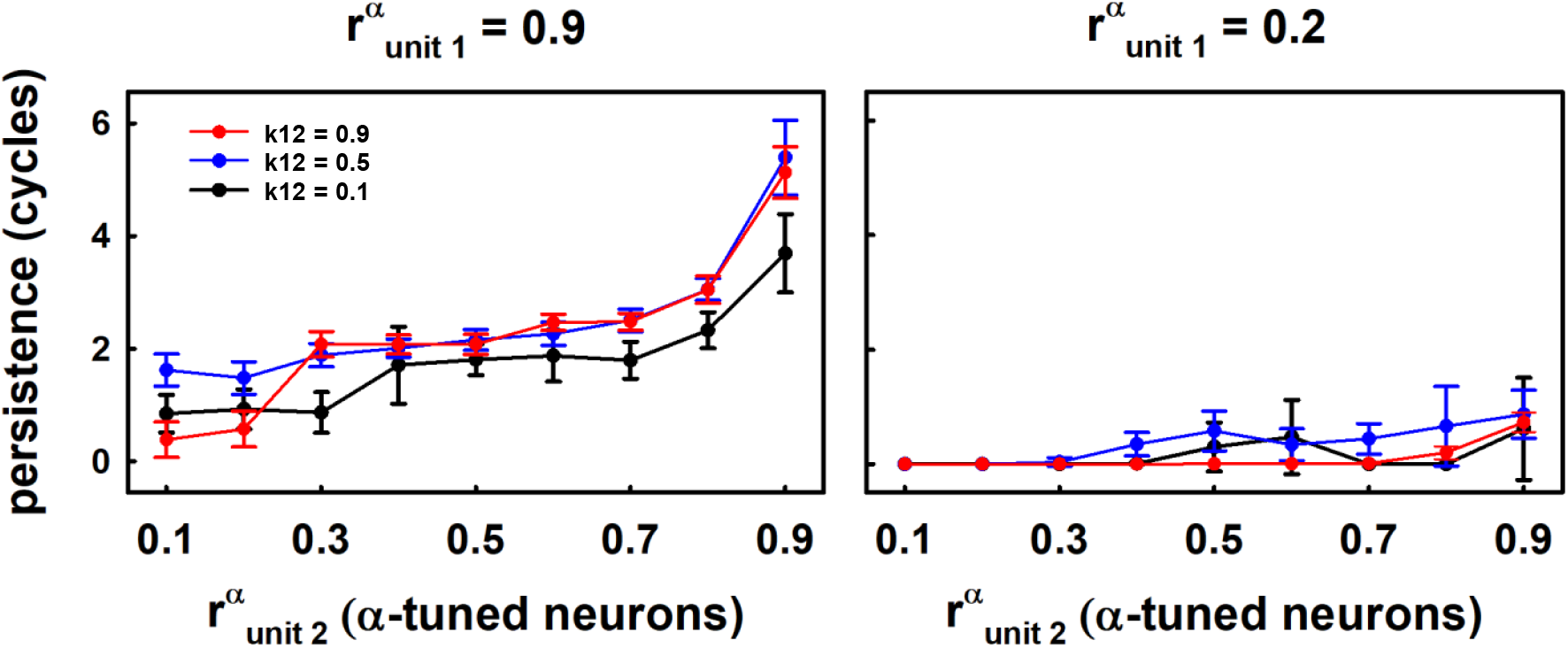
Effect of *α* proportion *r^α^* and coupling strength on persistence duration. Plots represent the mean and standard deviation of the persistence duration of the second column of the pair obtained for *k*12 = 0.1 (black lines), *k*12 = 0.5 (blue lines) and *k*12 = 0.9 (red lines). The first column only received the driving frequency and we present the case for *r^α^* = 0.9 **(left panel)**, and *r^α^* = 0.2 **(right panel)**

**Figure 11:**
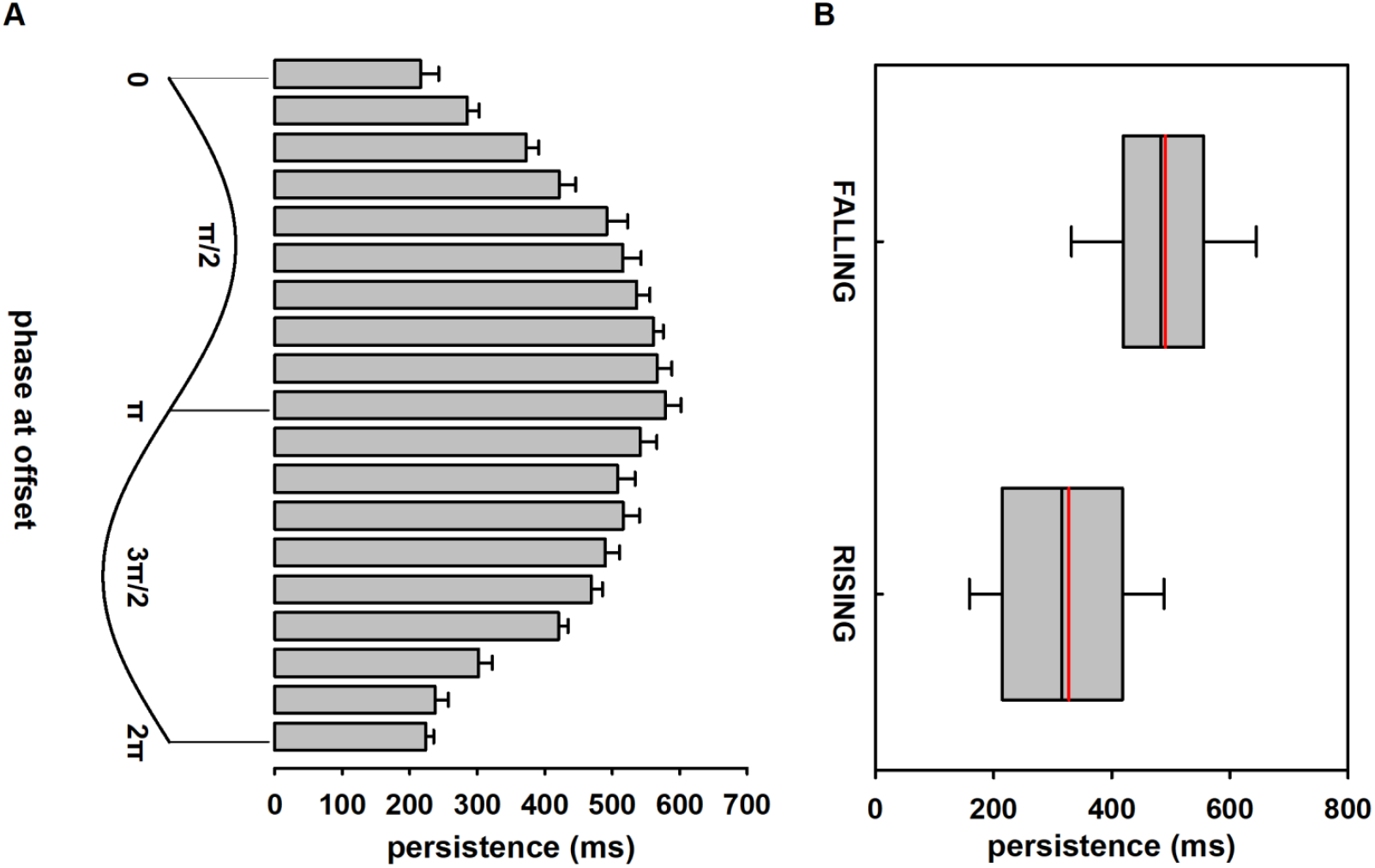
Persistence duration of the EEG like activity of a Jansen’s modified NMM as a function of the phase of the oscillation at the time the driving force was removed. (**A**) Persistence duration obtained for each of the eighteen sampled phases. Bars represent the mean and standard error calculated for N=30. (**B**) Falling and rising phases of the EEG-like activity (EEG-like phases towards the trough and the peaks of the alpha-oscillation, respectively) at the stimulus offset were grouped together, and plotted according to its PD. Each plot represents the mean (red vertical line) and median (black vertical line) of the PD computed for N = 270).

Therefore, our results provide insights about the anatomical limits of the propagation of entrainment, suggesting that the propagation will be restricted to cortical regions sharing similar intrinsic frequencies, hence processing the same type of information. This study confirms that Jansen-Rit based models are a suitable tool to study basic mechanisms of neural oscillations, including the persistence and propagation of neural entrainment. Future studies about the propagation of entrainment over large-scale cortical regions might involve the connection of a greater number of cortical Jansen’s columns, similar to that used to generate ongoing EEG activity in [39]. These investigations will contribute to advance the knowledge of the possible mechanisms underlying neural entrainment, relevant for the design of neuro-modulation strategies based on repetitive stimulation.

## Supplementary Materials

### Supplementary Tables

**Table 1:** Parameters used for the post-synaptic potential (PSP) generation of the *α* population

| Parameter | Biophysical Interpretation | Standard Value |
| --- | --- | --- |
| A | Maximum amplitude of the excitatory population | 3.25 mV |
| a | Lumped parameter: Spatially distributed delays | 100 $s^{-1}$ |
| B | Maximum amplitude of the inhibitory population | 22 mV |
| b | Lumped parameter: Spatially distributed delays | 50 $s^{-1}$ |

**Table 2:** Parameters used for the post-synaptic potential (PSP) generation of the *γ* population.

| Parameter | Biophysical Interpretation | Standard Value |
| --- | --- | --- |
| A | Maximum amplitude of the excitatory population | 11.375 mV |
| a | Lumped parameter: Spatially distributed delays | 350 $s^{-1}$ |
| B | Maximum amplitude of the inhibitory population | 132 mV |
| b | Lumped parameter: Spatially distributed delays | 300 $s^{-1}$ |

**Table 3:** General parameters of the connectivity constants between sub-populations of the model

| Significance | Parameter | Standard Value |
| --- | --- | --- |
| General constant | $C$ | 135 |
| Synapses between pyramidal cells and excitatory interneurons | $C_1$<br>$C_2$ | C<br>0.8*C |
| Synapses between pyramidal cells and inhibitory inter-neurons | $C_3$<br>$C_4$ | 0.25*C<br>0.25*C |

**Table 4:** General parameters of the Sigm function of the model

| Parameter | Biophysical Interpretation | Standard Value |
| --- | --- | --- |
| $e_0$ | Maximum firing rate | 2.5 |
| $v_0$ | Firing threshold | 6 mV |
| $s$ | Steepness of the sigmoidal function | 0.56 |

### Supplementary Equations

System of equations for the alpha population for both columns. These equations are equivalent for the *γ* population.

The outputs of the three postsynaptic boxes in the first column were noted as: the resulting from the pyramidal cells *y*0, the output of the excitatory inter-neurons *y*1 and the output of the inhibitory inter-neurons *y*2. After coupling the columns, the output of these three postsynaptic boxes, for the second column, were termed as *y*6, *y*7 and *y*8. Furthermore, each postsynaptic box was computed having into account that they are composed by two populations (*α* and *γ*) and the proportion that represent each of this populations in every column (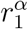 and 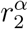).

Thus, *y*0 and *y*6, representing the outputs of the postsynaptic boxes of the columns 1 and 2 respectively, were computed as:

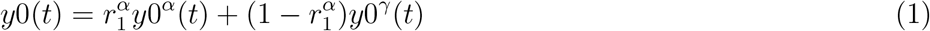

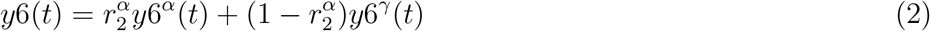

An equivalent reasoning is made for computing the output of the excitatory branch for both columns: *y*1(*t*) for column 1, and *y*7(*t*) for column 2. The resulting equations are given by:

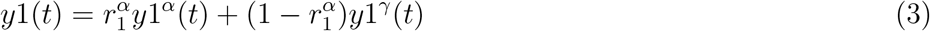

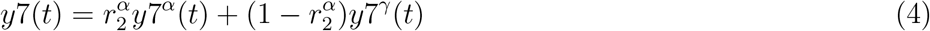

The output for the inhibitory branch for columns 1 and 2 are *y*2 and *y*8 respectively, computed as:

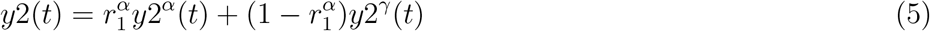

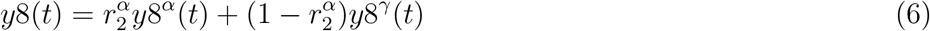

Furthermore, the postsynaptic membrane potential in the pyramidal cell (equivalent to the EEG *y*(*t*) = *y*1(*t*) − *y*2(*t*)), were computed for each column in the coupled NMM. They were noted as *y*_*C*1_(*t*) and *y*_*C*2_(*t*) for columns 1 and 2, respectively, and computed using equations 1–6 as follows:

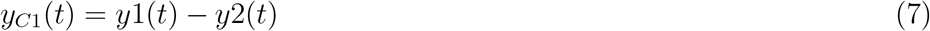

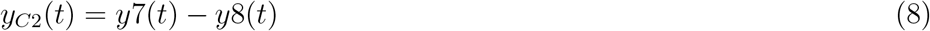

The system of equations of the model, is composed by the system of equations describing the *α* population and the system of equations describing the *γ* population for every column. As a matter of example, we described the system of equations for the alpha population in both columns. Nevertheless, all the codes from the model are available as Supplementary Materials.

#### Alpha population for the first column

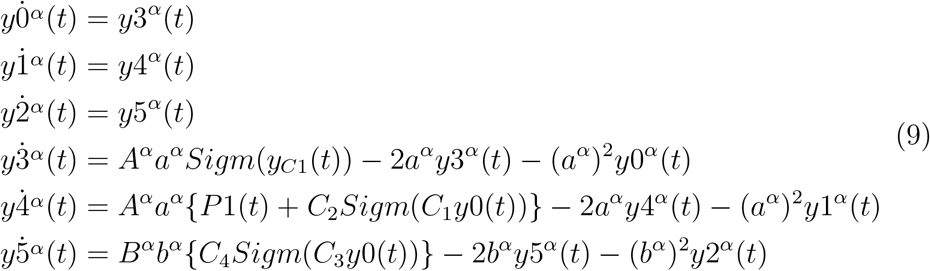

#### Alpha population for the second column

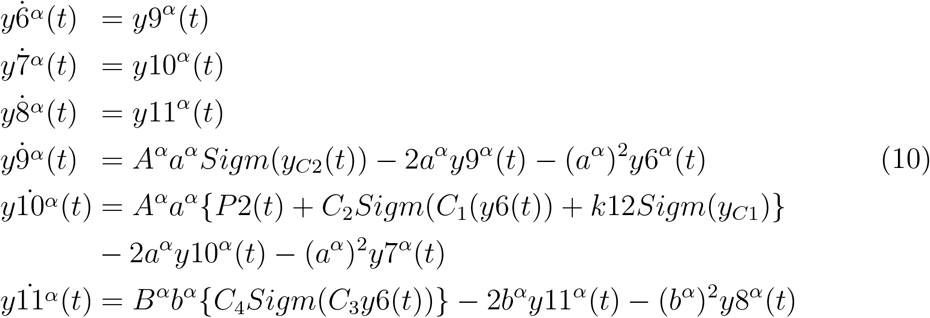

### Supplementary Results

#### Initial Simulations

Intrinsic Oscillations depends on the *α* proportion parameter *r^α^*.

We analyzed intrinsic oscillations generated with a single column NMM, where the proportion of the *α* proportion was varied between *r^α^* = 0 and *r^α^* = 1 in steps of 0.01. Fifty simulations (trials) were run for each value of *r^α^*, in which each simulation consisted of 1000 time steps (1s). For each trial, the power spectrum was calculated using the discrete Fourier transform and the mean power spectrum was computed for each *r^α^* value. Trials of the same *r^α^* value were averaged in the time-domain to obtain the time series presented in Supplementary Figure 1. The extrinsic input was sampled from a Gaussian distributed noise, which represents a pulse density uniformly distributed between 120 and 320 spikes per second, modeling the background noise in the cerebral cortex.

Supplementary Figure 1 presents examples of the time series and the power of intrinsic oscillations for different values of *r^α^*. We can observe two examples of gamma oscillations corresponding to *r^α^* = 0.2 and *r^α^* = 0.3, and alpha oscillations using *r^α^* = 0.8 and *r^α^* = 0.9 ( [35,38]). A prominent feature of the NMM was the generation of oscillatory activity to a noise input, which was highly dependent on *r^α^*. While the power spectrum of the neuronal activity was dominated by gamma oscillations when *r^α^* was relatively small (*r^α^* = 0.2, and *r^α^* = 0.1), alpha oscillations resulted when the noise input was presented to NMM with (*r^α^* = 0.8, and *r^α^* = 0.9). For NMM with *r^α^* = 0.2, and *r^α^* = 0.9, the peak amplitude was obtained at 43 and 11 Hz, respectively. Slight shifts of the peak frequency of the intrinsic oscillations toward the middle region of the power spectrum were observed when *r^α^* changed to *r^α^* = 0.3 compared to *r^α^* = 0.2 and when *r^α^* value was *r^α^* = 0.8 compared to *r^α^* = 0.9. These shifts in the peak frequency were accompanied by oscillations with broader frequency spectrum. The power of gamma oscillations was higher than that of the alpha oscillations, when NMM with equivalent *r^α^* are compared (*r^α^* = 0.1 against *r^α^* = 0.9, for example). This result can be explained by the different post-synaptic potentials (PSP) functions of the excitatory and inhibitory neurons comprising the *α* and *γ* populations (Supplementary Figure 3).

**Figure 1:**
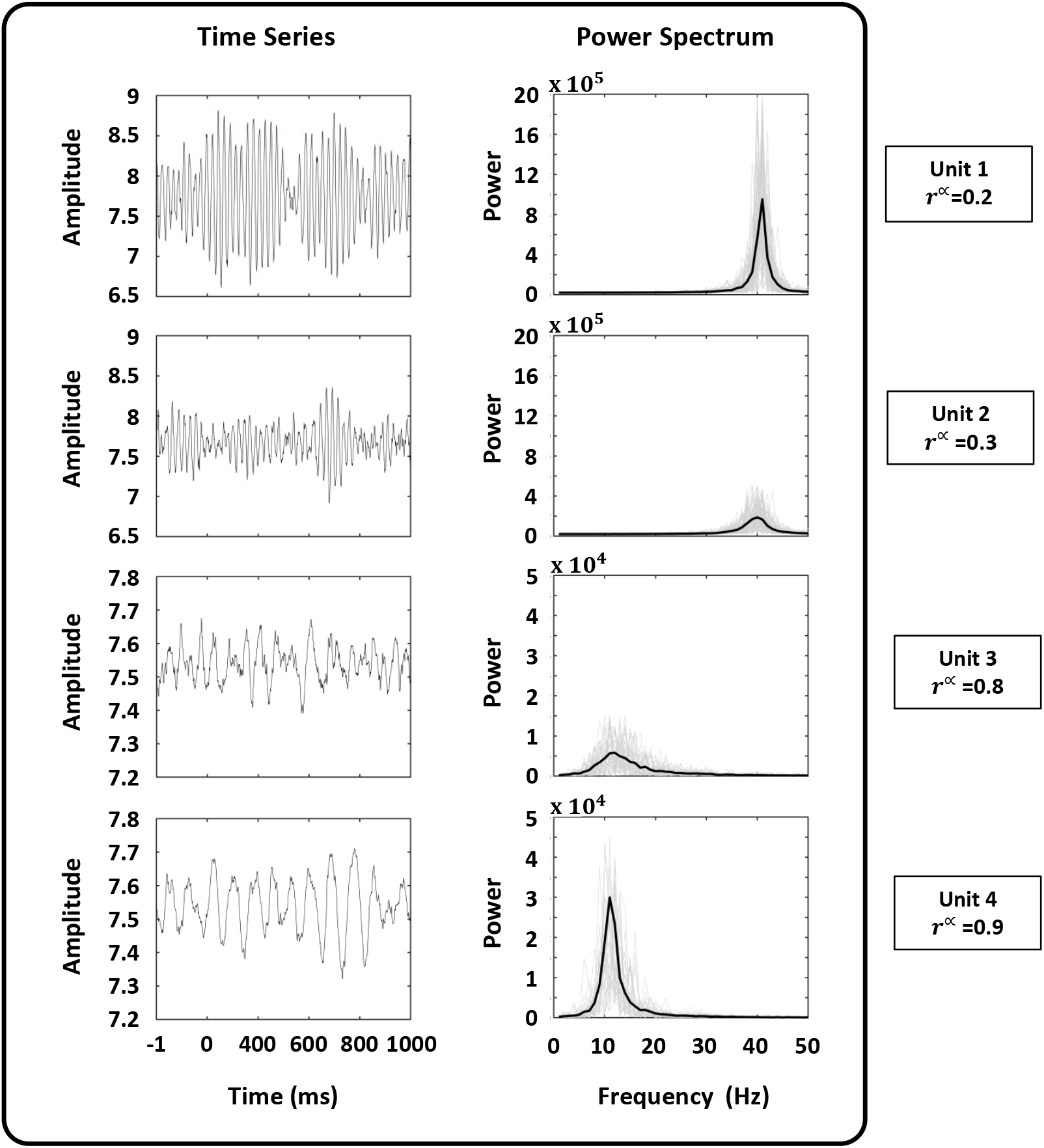
Examples of intrinsic oscillations of single independent units. Time series and Power Spectrum of independent units obtained varying the parameter of the model *r^α^* are shown. Using *r^α^* = 0.2 and *r^α^* = 0.3 intrinsic oscillations in the gamma range are obtained, while with *r^α^* = 0.8 and *r^α^* = 0.9 alpha oscillations emerged.

A generalization of the effect of *α* proportion *r^α^* on the spectral profile of the NMM is presented in Figure 2. Simulations were run such that *r^α^* was varied between 0 and 1, in 0.1 steps. The spectral profile of all simulation (50 repetitions for each *r^α^*) are plotted together. The results confirmed those presented in Supplementary Figure 1, and extended the findings to populations with other relative proportion of *α*-tuned neurons (*r^α^*).

The spectral profile of the single-column two-populations NMM was characterized by peak amplitudes of alpha and gamma oscillations when *r^α^* was close to 0 and 1, respectively. In those cases, the NMM was tuned to generate *α* or *γ* oscillations for the noise inputs implemented in this study. Consequently, the output of the model was extremely consistent across repetitions (runs of the simulations) for *r^α^* close to 0 and 1. This was reflected in the narrow spectral profile of the oscillatory activity obtained at those *r^α^*. The generation of alpha oscillations expanded for 0.8 ≤*r^α^* ≤ 1, while the generation of gamma oscillations expanded for 0 ≤ *r^α^* ≤ 0.3. The spectral profile obtained for intermediate values of *r^α^* were a consequence of having NMM with different mixes (combinations) of *α* and *γ*-tuned neural populations. As expected, these mixed neuronal ensembles made the oscillatory behavior of NMM with intermediate *r^α^* display a wider frequency profile than those presented by NMM with extreme values of *r^α^*.

**Figure 2:**
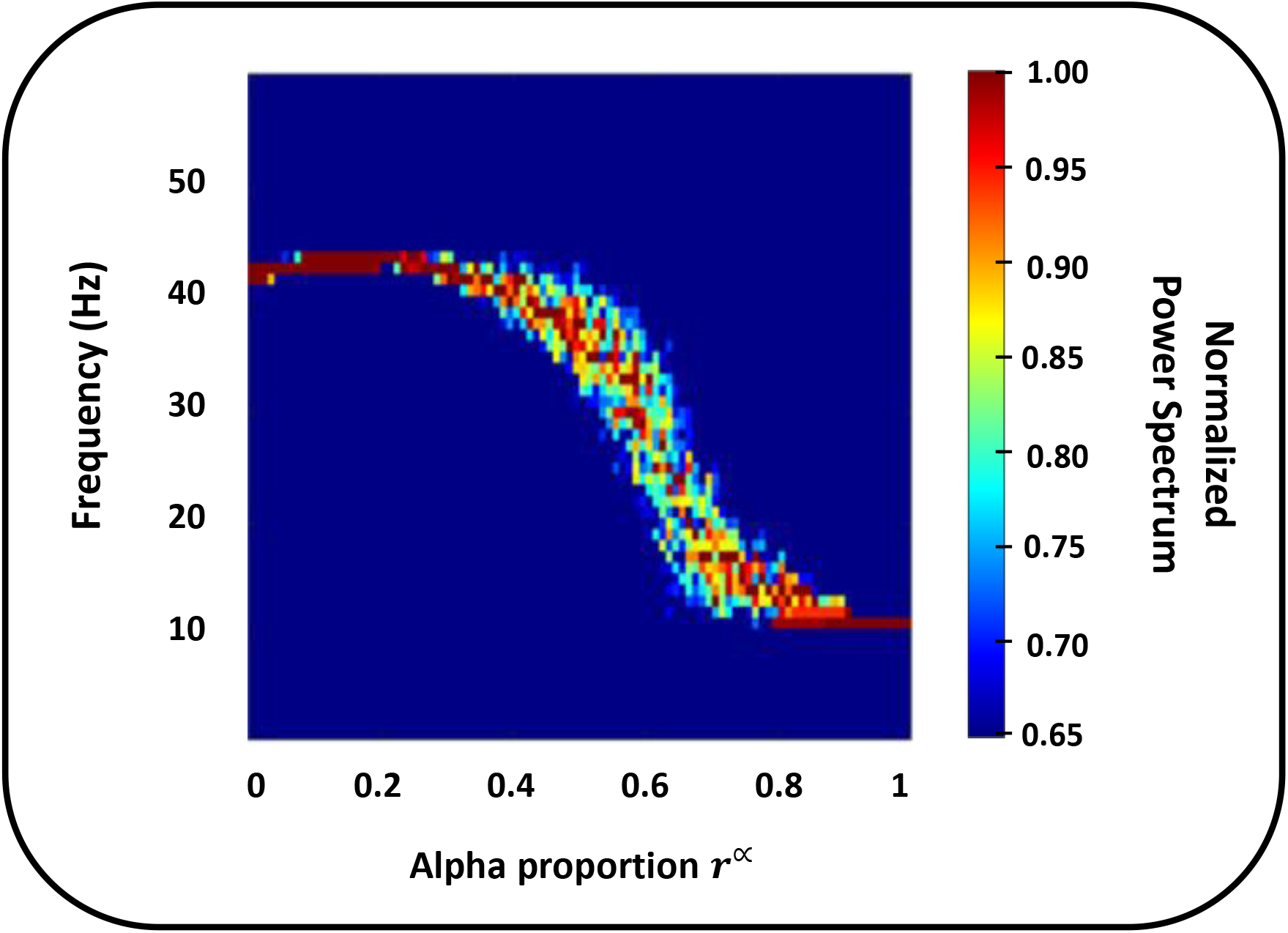
Spectral representation of the intrinsic oscillations of the model as a function of the *α* proportion *r^α^*.

#### Comparing Excitatory and Inhibitory PSPs for *α* and *γ* populations

The PSP functions (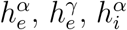 and 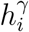,) for both *α* and *γ* populations, composed by excitatory and inhibitory sub-populations are presented in Figure 3. It is noteworthy that temporal dynamics are different between *α* and *γ* populations due to their PSP functions. Slower temporal evolution and smaller amplitude are found in *α* population PSPs in comparison to those presented by the *γ* population. At the same time, there is a difference between inhibitory PSP (IPSP) and excitatory PSP (EPSP) of any particular population (*α* and *γ*). IPSP presented increased amplitude and longer duration as compared to the corresponding EPSP. These behaviours are due to parameters *A^α,γ^*, *B^α,γ^* and *a^α,γ^* and *b^α,γ^* of the equations, which control the PSP functions amplitudes and temporal courses.

**Figure 3:**
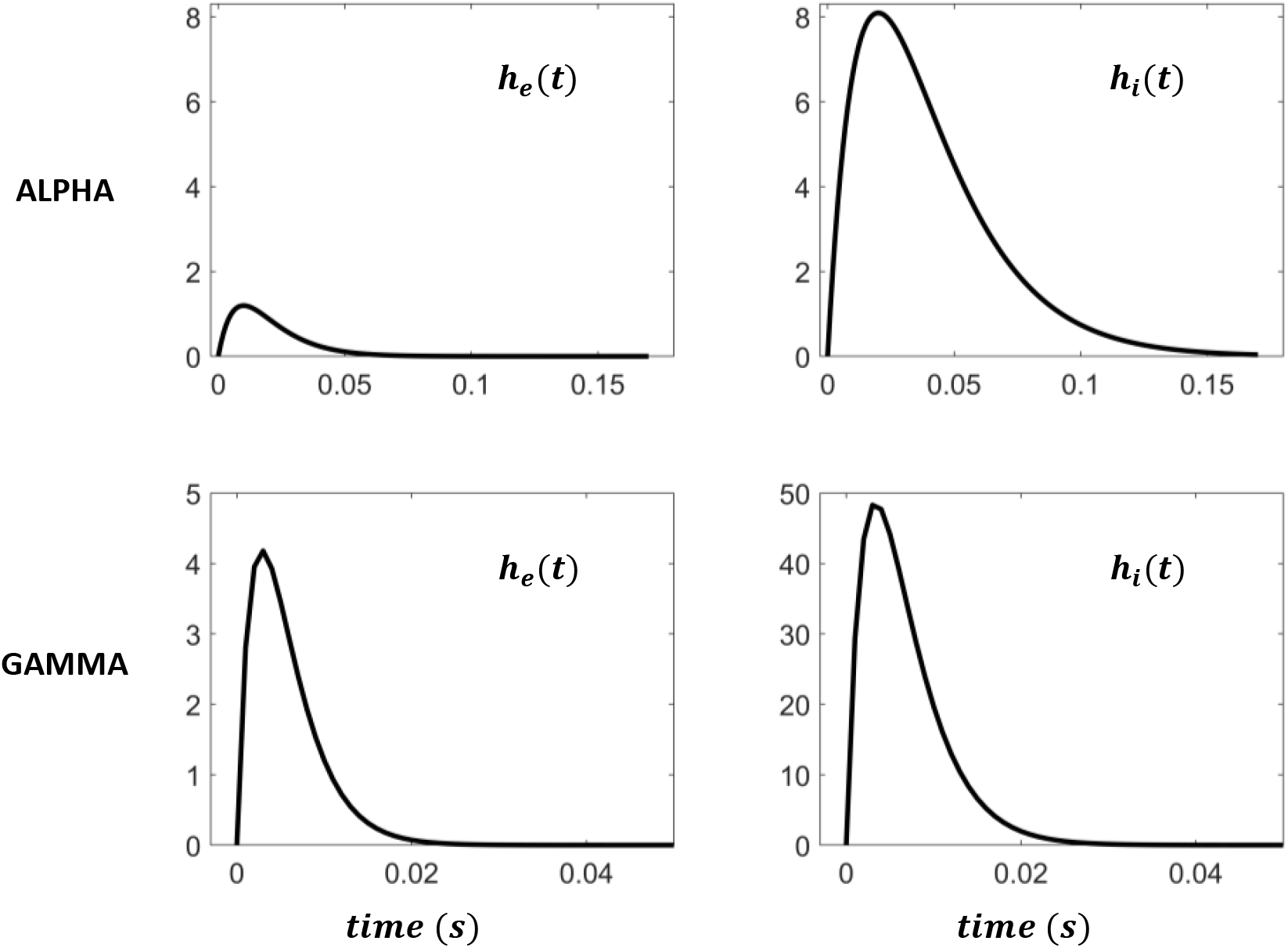
Comparison between the Excitatory and Inhibitory Postsynaptic Potentials generating functions for *α* and *γ* populations. **(Top panels)** Dynamics of excitatory (left chart) and inhibitory PSP (right chart) of *α*-tuned sub-populations (Parameters in Table 1). **(Lower panels)** Equivalent *γ* dynamics are shown (Parameters in Table 2).

